# Microbial Crosstalk in the Neonatal Gut: *Candida albicans* Impacts Group B *Streptococcus* Pathogenicity

**DOI:** 10.64898/2026.10.04.756596

**Authors:** April K. Lindon, Alexia N. Pearah, Kristen Dominguez, Nichol John-Lewis Edwards, Oriana M. Williams, Ojas Natarajan, Ryan Handel, Cinthya N. Zelaya Aparicio, Larry J. Dishaw, Sophie E. Darch, Tara M. Randis

**Affiliations:** Pediatrics, Morsani College of Medicine, University of South Florida, Tampa, FL; Molecular Medicine, Morsani College of Medicine, University of South Florida, Tampa, FL

## Abstract

**Background:** Group B *Streptococcus* (GBS) is a leading cause of bacterial sepsis globally. Late-onset (LO) GBS disease is preceded by colonization within the gastrointestinal tract in vulnerable neonates. Previous studies demonstrate that *Candida albicans* (*C. albicans*) promotes GBS colonization of host tissues including the bladder and vagina. However, its impact on GBS colonization of the intestinal tract is unknown.

**Methods:** We established a new animal model to study GBS and *C. albicans* co-colonization model in the context of the newborn host to examine *C. albicans* impact on GBS and further evaluated fluconazole prophylactic treatment to these same outcomes. Physical interactions between organisms *in vivo* were imaged and GBS gene expression was evaluated.

**Results:** *C. albicans* increased GBS burden in the neonatal gut and GBS was physically colocalized with *C. albicans* hyphae as evidenced by the colocalization analysis. Fluconazole treatment of co-colonized mice decreased extraintestinal GBS dissemination. *C. albicans* presence upregulated genes encoding proteins responsible for GBS adhesion and invasion of intestinal epithelial cells.

**Conclusion:** *C. albicans* enhances GBS burden and pathogenicity in the neonatal gut. These findings highlight the need for further investigation into *C. albicans* as a potential risk factor for GBS colonization and LO disease in human infants.

## Introduction

Group B *Streptococcus* (GBS) is a leading cause of bacterial infection and death of newborns.^1^ Global estimates of burden exceed 100,000 cases annually.^2^ Mortality due to late-onset (LO) GBS is ∼5% overall, though notably higher (>20%) in low-birth-weight infants, who are particularly susceptible.^3–5^ LO GBS infection, presenting beyond the first week of life, is now the most common clinical manifestation of invasive GBS disease during infancy in the US and other regions that have adopted the strategy of intrapartum antibiotic prophylaxis for individuals who are colonized with GBS during pregnancy.^6^

LO GBS infection occurs when GBS colonizing the gastrointestinal tract crosses intestinal barriers to cause invasive disease.^7,8^ It typically manifests as sepsis, bone/soft tissue infections, or meningitis, all of which are associated with significant long-term sequelae in survivors.^2^ What drives the progression from GI colonization to invasive LO disease is poorly understood and has slowed the development of preventive strategies in exposed infants.

*Candida albicans* (*C. albicans*) is another pathobiont that colonizes the intestinal tract in 30-60% of healthy newborns.^9,10^ Similar to GBS, intestinal colonization with *C. albicans* can lead to invasive infection in vulnerable hosts.^11,12^ Importantly, *C. albicans* has been shown to modulate the virulence of other bacteria, and synergistic interactions between *C. albicans* and bacteria such as *Staphylococcus aureus*,^13,14^ *Escherichia coli*,^15,16^ and *Pseudomonas aeruginosa*,^17,18^ are well described. *C. albicans* forms polymicrobial biofilms, by providing structural support, increasing bacterial adherence, and offering protection from host immune responses and antimicrobial agents.^19^ In addition, *C. albicans* hyphae may cause physical damage to host barriers, thus allowing other microorganisms to penetrate more easily.^20,21^ In the absence of some bacteria, such as Lactobacilli, *C. albicans* filamentation is less constrained. Thus, *C. albicans* virulence is linked to dysbiosis, which is quite common during infancy, especially in prematurity.^22–24^

Synergistic interactions between *C. albicans* and GBS occur at bladder and vaginal mucosal surfaces. *In vitro* studies reveal that *C. albicans* promotes GBS colonization of human vaginal epithelial cells, specifically via binding of a GBS polypeptide adhesin, BspA, with the *C. albicans* fungal hyphal-associated adhesin, Als3.^25^ In a mouse model of urinary tract infection, *C. albicans* enhances GBS colonization of the bladder epithelium, an interaction that similarly is dependent on the expression of Als3.^26^ Another recent study co-infected zebrafish with GBS and *C. albicans* revealing increased GBS burden in the co-infected model versus the GBS monocolonized model.^27^ Despite growing recognition of polymicrobial interactions in mucosal colonization, studies examining the role of *C. albicans* in modulating GBS activity have been limited to the vaginal and bladder epithelia. The objective of this study was to determine the impact of *C. albicans* on GBS intestinal colonization and progression to invasive disease using a newly established neonatal mouse model of co-colonization.

## Methods

### Microbial Cultures and Conditions

GBS COH-1 (serotype III ST-17, BAA-1176™) was grown overnight at 37°C in tryptic soy broth (TS) and then enumerated by completing serial dilutions on CHROMagar™ StrepB plates. *C. albicans* (Robin) Berkhout strain ATCC 10231 was incubated in yeast malt broth at 37°C for 24 hours under continuous shaking at 200 rpm for animal experiments and grown stationary for gene expression and cell experiments. Inoculum was enumerated by completing serial dilutions on CHROMagar™ Candida.

### Animal Experiments

#### GBS-*C. albicans* co-colonization model

All experiments were approved by and performed in accordance with the University of South Florida Institutional Animal Care and Use Committee. We used our previously published post-natal GBS colonization model^8^ with the following modifications to establish a novel GBS-*C. albicans* co-colonization model. C57BL6/J breeder pairs were purchased from Jackson Laboratories and after acclimating, mice were mated and pregnant females were housed separately for the remainder of the experiment. Pups were infected on day of life 7 with 50 µl of *C. albicans* suspension via oral gavage (stationary phase culture was pelleted and resuspended in sterile phosphate-buffered saline (PBS) to a final concentration of 10^8^ CFU/mL for dose of 5 X 10^6^ CFU). On days of life 10 and 11, pups were infected with 100 µl of GBS (stationary phase culture was pelleted and resuspended in sterile PBS to a final concentration of 10^8^ CFU/mL for final dose of 10^7^ CFU daily) (Fig. 1a). For GBS monoinfection, 50 µl of PBS vehicle control was administered on Day of life 7 instead of *C. albicans* (Fig. 1b). All littermates remained with their biologic dam for the remainder of the experiment. Animals were monitored daily and weighed 3 times per week to assess long-term mortality. In a separate cohort, mice co-infected with GBS and *C. albicans* received fluconazole (Sigma F8929, 5mg/kg daily, intraperitoneal) or vehicle control administered daily from days of life 11 to 14.

**Figure 1.**
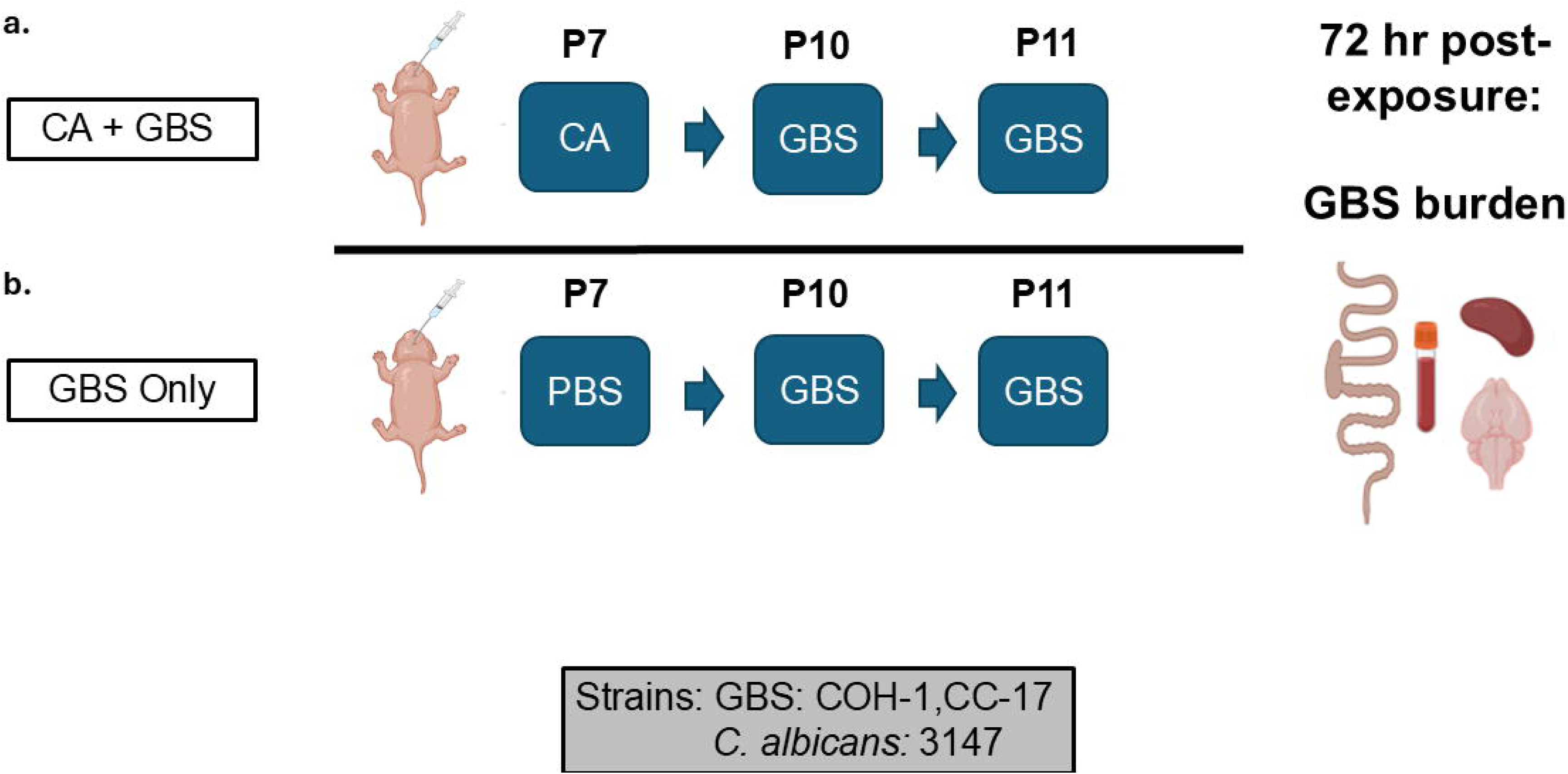
GBS and *C. albicans* co-colonization LO disease model. GBS and CA co-infection (**a**) was accomplished by infecting with CA at a dose of 5 X 10^6^ CFU on Day of life 7 followed by 2 doses of GBS on Day of life 10 and 11 with a final dose of 10^7^ CFU daily. GBS monoinfection (**b**) 50 µl of PBS was administered by oral gavage on Day of life 7 instead of CA. GBS and CA burden was evaluated by completing serial dilutions 72 hours post exposure of the small intestine and colon as well as extraintestinal dissemination to the blood, brain and spleen. P = Postnatal day (P7 = Postnatal day 7)

### Tissue processing

Mice were euthanized on day of life 14, and blood, spleen, brain, and intestinal tissues were extracted to determine GBS burden. Tissues were collected in PBS and gently homogenized using the Bullet Blender (Next Advance Inc.) for 30□seconds, speed 3. Serial dilutions of tissue homogenates were plated on selective media (CHROMagar^TM^ StrepB) to determine GBS burden in tissue (reported as CFU per gram tissue) and blood (100□μl plated). To evaluate *C. albicans* burden, serial dilutions were plated and enumerated on selective media (CHROMagar^TM^ Candida).

### Murine fixed tissue staining

Pups were co-infected as described above and following euthanasia, small intestine and colon were fixed in 4% paraformaldehyde (PFA) for 24 hours, then transferred to a 70% ethanol solution prior to being embedded in paraffin for sectioning. Fixed murine tissue slides were stained using the following protocol adapted from Abcam.^28^ Deparaffinization was completed by washing with Safe Clear^TM^ and ethanol in the following 3 minute increments with gentle agitation: 2 washes with Safe Clear; 1 wash containing Safe Clear 1:1 with 100% ethanol, 2 washes with 100% ethanol, 1 wash with 95% ethanol, 1 wash with 70% ethanol, 1 wash with 50% ethanol. Slides were rinsed with cold tap water and then immersed in a Coplin jar containing sodium citrate buffer pH 6.0 and placed in a 70°C water bath overnight for antigen retrieval. The following day, slides were permeabilized by immersing in PBS with 0.5% Tween20 for 10 minutes, followed by staining as follows: slides were washed 2 x for 5 minutes in PBS containing 0.025% Triton x-100 while under gentle agitation and blocked in 10% goat serum with 1% BSA in PBS for 1 hour. Slides were incubated with GBS primary antibody (Abcam: ab53584; rabbit polyclonal Group B Streptococcus antibody, 1:200 dilution in 1% BSA in TBS) and *C. albicans* antibody (Abcam: ab21164; rabbit polyclonal *Candida albicans* antibody conjugated to FITC, 1:200 dilution in 1% BSA in TBS) for 1 hour at room temperature. Following incubation, slides were washed 2 times for 5 minutes with PBS containing 0.025% Triton x-100 under gentle agitation. Fixed tissue slides were incubated with secondary antibody Alexa Fluor 568 Anti-rabbit (1:500 dilution in 1% BSA in TBS) for 1 hour at room temperature, washed 3 times in 5-minute increments with PBS, and coverslip-mounted using ProLong^TM^ Diamond Antifade Mountant with DAPI (Invitrogen^TM^ P36966). Samples were stored at 4°C until imaging.

### Murine stained fixed tissue imaging and colocalization analysis

Murine colonic fixed and stained tissue samples from four co-colonized mice (across three experimental litters) were examined, with four distinct hyphae-containing tissue sections per specimen rendered and analyzed (16 total images). The Zeiss Laser Scanning Confocal Microscope 880 was used to capture z-stacked images collected at 0.5 µm intervals with 100× oil objective (NA 1.4). To identify the presence and absence of staining performed on fixed murine tissue, we used a 2-channel approach: Alexa Fluor 568 (excitation at 578 nm, emission at ∼ 603 nm) for GBS and FITC (excitation at 491 nm, emission at ∼ 516 nm) for *C. albicans*. Images were subsequently uploaded into Imaris (v10.2, Oxford Instruments) and a colocalization analysis of GBS proximity to *C. albicans* was completed as previously established by Williams & Darch, 2026 with minor adaptations.^29^ Isosurfaces were assigned to voxels of Alexa Fluor 568 stained GBS and separately to the voxels of FITC stained *C. albicans*. GBS isosurfaces were filtered by distance to *C. albicans* isosurfaces. The GBS isosurfaces that had a distance of zero to *C. albicans* isosurfaces were selected and a new duplicate isosurface of the GBS stain was created to represent the overlap with *C. albicans*. Percent overlap was calculated from the GBS voxel overlapped with *C. albicans* by the total GBS voxel volume.

### Gene expression

GBS and *C. albicans* were grown overnight as described previously followed by diluting 1:100 and incubating for 4 hours to mid-logarithmic phase. RNA was extracted from the *in vitro* GBS monocultures and GBS - *C. albicans* co-cultures using the Zymo Research Corporation Quick-RNA Fungal/Bacterial MiniPrep™ Kit (Zymo Research R2014) according to the manufacturer’s protocol, followed by Invitrogen™ Ambion™ DNase I (RNase-free) treatment. The concentration and purity of the RNA were measured using the NanoDrop One with the A260/280 ratio confirmed to be within 1.8–2.0 for all samples. cDNA was synthesized using the High-Capacity cDNA Reverse Transcription Kit (Applied Biosystems) following the manufacturer’s instructions. A control reaction without reverse transcriptase (No-RT control) was included for each sample to confirm the absence of genomic DNA contamination. Quantitative real-time PCR was performed using SYBR Green Supermix (Bio-Rad) on Bio-Rad CFX Manager. The thermal cycling protocol was as follows: polymerase activation and DNA denaturation at 95°C for 30 seconds, followed by 40 cycles of 95°C for 10 seconds and 60°C for 30 seconds. Gene-specific primers for the target genes *cylE*, *hvgA*, *srr2*, *fbsB*, and *spb1* were used. Primer sequences were previously published and are listed in Table S1. Technical replicates (3) were run for each biological sample (4), and a no-template control (NTC) was included for each primer pair on every plate. The Ct values of the target genes were normalized to the reference gene *recA* as the endogenous control and relative quantification was performed using the ΔΔ^-CT^ method. The GBS monoculture was used as the baseline reference, with its expression level normalized to 1.0 for log□ fold change calculations. All plotted data represent the log₂ fold changes in expression of each target gene relative to this baseline. Experiments were repeated a minimum of 3 times to ensure reproducibility.

### Cell culture

Caco-2 (ATCC HTB-37) and T-84 (ATCC CCL-24) cell lines were used in all cell culture experiments. Caco-2 cells were cultured in Eagle’s minimum essential medium (EMEM, Corning 10-009-CV) supplemented with 10% fetal bovine serum (FBS, Gibco 10082-147) and 1% penicillin–streptomycin (Gibco 15140-122). T-84 cells were cultured in Dulbecco’s minimum essential medium/F-12 (DMEM/F-12, Corning 10-090-CVR) supplemented with 5% FBS and 1% penicillin–streptomycin. Cells were grown to 80% confluence on 24-well polystyrene plates (Falcon 353047) and media was changed every 3–4 days.

### Adhesion and invasion assay

GBS was grown to mid-logarithmic phase in TS broth as noted above, centrifuged for 7 minutes at 3,000 g and resuspended in EMEM 10% FBS for Caco-2 cell experiments and resuspended in DMEM/F-12 5% FBS for T84 cell experiments to obtain a final concentration of 10^8^ CFU/mL to achieve a multiplicity of infection (MOI) of 50. *C. albicans* was cultivated in yeast malt broth as mentioned above, centrifuged for 7 minutes at 3,000 g and resuspended in either EMEM or DMEM for cell culture experiments. The same infection concentration was used for *C. albicans* as GBS.

Caco-2 cells were seeded into a 24-well plate and incubated at 37°C and 5% CO_2_ until they reached 80% confluency. To model co-exposure, cells were first incubated with *C. albicans* for 2 hours, supernatant removed and followed by a 2-hour incubation with GBS. Cells exposed to only GBS were incubated for 2 hours with the COH-1 strain.

For the adhesion assay, after the incubation period, media was aspirated from the wells and cells were washed with PBS three times. Cells were disassociated from the plate using TripLE^TM^ (Gibco^TM^ 12-604-013) and mechanical disruption with antibiotic-free media. Each well was placed into its own respective Eppendorf tube and serial dilutions plated on CHROMagar^TM^ Strep B plates and CHROMagar^TM^ Candida plates to the 9^th^. CFU was completed for COH-1 the next day and CFU for *C. albicans* CFU were completed 2 days after plating.

For the invasion assay, following GBS/CA exposure, media was aspirated from the wells and cells were washed with PBS three times. Cells were incubated for 2 hours with EMEM or DMEM containing 100 µg/mL of gentamicin and 5 µg/mL of penicillin/streptomycin to remove extracellular bacteria and washed 3 times with PBS. Cells were lysed with 0.1% Triton x-100 in PBS for 30 minutes, TripLE^TM^ for dissociation from the plate, then the contents of each well were placed into a separate Eppendorf tube. Serial dilutions were completed as previously described with the adhesion assay.

## Results

### *C. albicans* increased GBS colonization density

We successfully established a reproducible model of GBS and *C. albicans* co-colonization in the gastrointestinal tract of neonatal mice (Fig. 1). Mice co-colonized with *C. albicans* and GBS had a significantly greater burden of GBS recovered from both the small intestine and the colon (Fig. 2a, b). Extraintestinal dissemination to remote organs (Fig. 2c) and mortality (Fig. 2d) were assessed in both groups, but no statistically significant difference was observed between groups. Of note, the increased GBS intestinal burden observed in the co-colonization model persisted regardless of the sequences of GBS or *C. albicans* exposure (Fig. S1).

**Figure 2.**
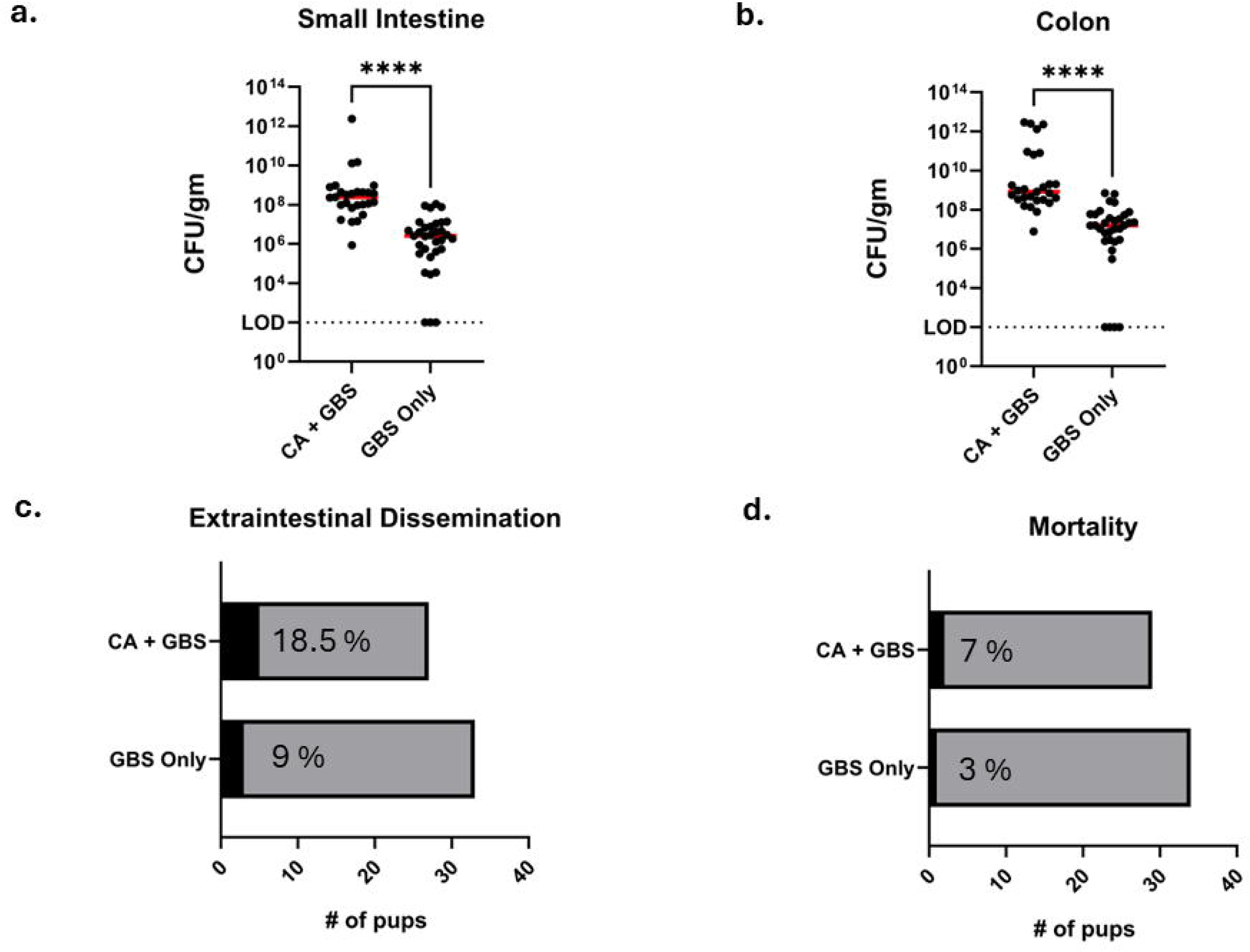
*C. albicans* impact on GBS colonization and dissemination *in vivo*. GBS Burden in small intestine (CA + GBS median CFU = 3.2 X 10^8^ CFU/gm tissue versus GBS monoinfection median CFU = 4 X 10^6^ CFU/gm tissue) (**a**) and colon (CA + GBS median CFU = 5.75 X 10^8^ CFU/gm tissue versus GBS monoinfection median CFU = 1.6 X 10^7^ CFU/gm tissue) (**b**) of co-colonized versus mono-colonized mice [****P<0.0001, Mann-Whitney test; Limit of detection=10^2^]. Extraintestinal dissemination (**c**) to the blood, brain, and/or spleen as well as mortality (**d**) were evaluated with no statistical significant difference between groups [Fisher’s test].

### Fluconazole treatment reduced extraintestinal GBS dissemination

Administration of fluconazole to mice co-colonized with *C. albicans* and GBS did not impact the overall tissue burden of GBS in small intestine or colon (Fig. 3a, b), however, fluconazole treatment significantly reduced the burden of *C. albicans* recovered from intestinal tissues (Fig. S3). There was a significant decrease in extraintestinal dissemination of GBS in animals co-colonized and treated with fluconazole versus those untreated (Fig. 3c). It is notable that dissemination of GBS in both treated and untreated study groups was higher than what was observed at baseline in our model of co-colonization, however, this may be attributed to the repeated handling of pups and stress of injections (PBS or drug) in both groups. There was no observed mortality in either group. Pups infected with *C. albicans* alone and treated with fluconazole exhibited complete clearance of fungal burden in the small intestine and a marked reduction in colonic burden. In contrast, fluconazole-treated co-colonized pups (*C. albicans* + GBS) showed a significant decrease in *C. albicans* burden relative to untreated co-colonized controls, yet they demonstrated a trend toward greater fungal persistence compared with the fluconazole-treated monoinfection group.

**Figure 3.**
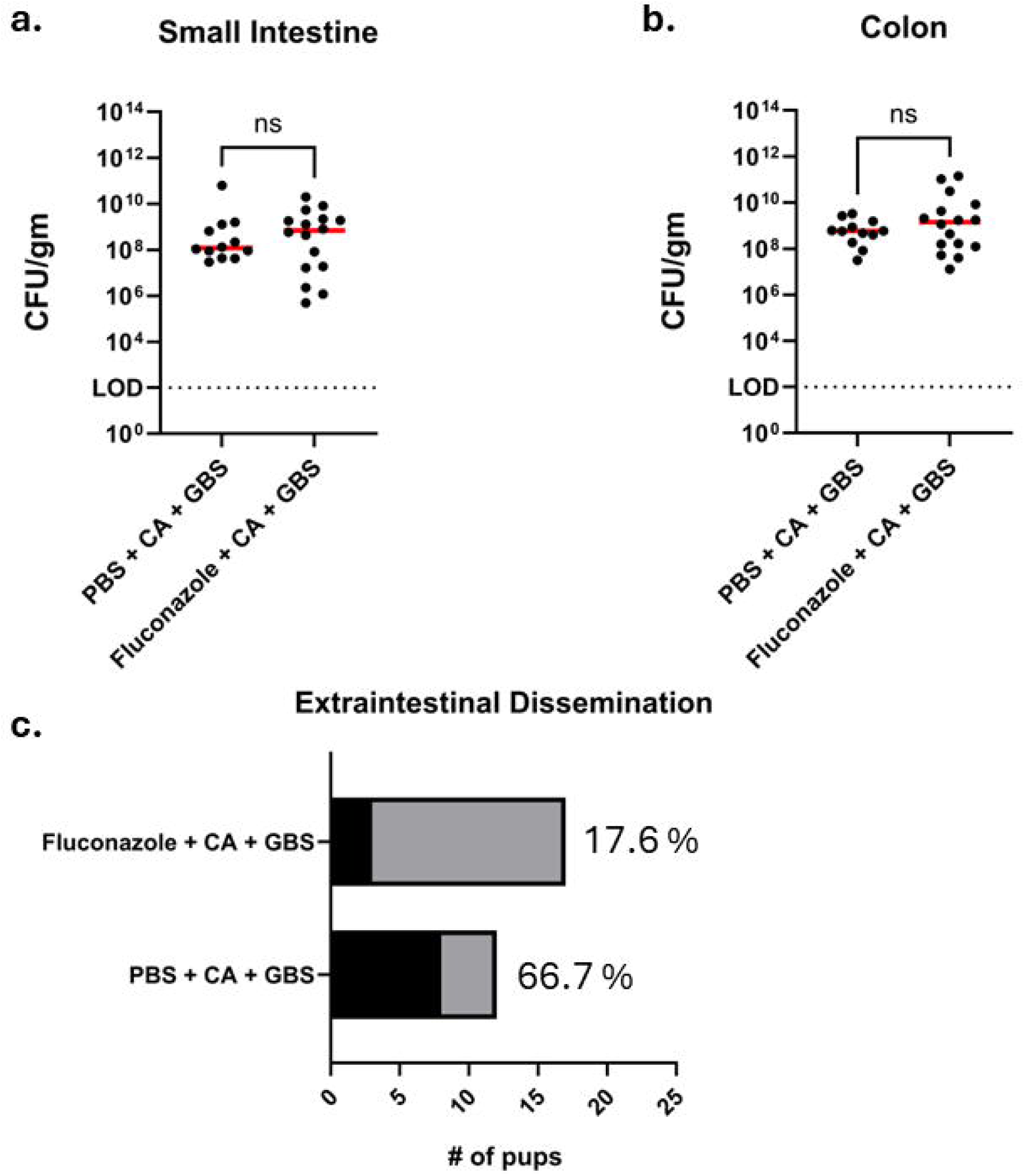
Mice Co-infected with GBS and *Candida albicans* treated with PBS vs Fluconazole. GBS burden in the small intestine (**a**) and colon (**b**) was evaluated [ns=nonsignificant, Mann-Whitney test]. Extraintestinal dissemination was also measured (**c**) with no mortality observed in either group [P=0.0177, NS=nonsignificant, Fisher’s exact test]. Limit of detection is 10^2^.

### GBS physically associates with *C. albicans* hyphae in the gut

Scanning confocal laser microscopy revealed GBS adhering to *C. albicans* hyphae in the colon (Fig. 4a, b, c). Signal from fluorescently labeled GBS and *C. albicans* overlapped spatially indicating co-localization. Spatial analysis demonstrated greater than 70 percent association of GBS to *C. albicans* across replicates (Fig. 4d).

**Figure 4.**
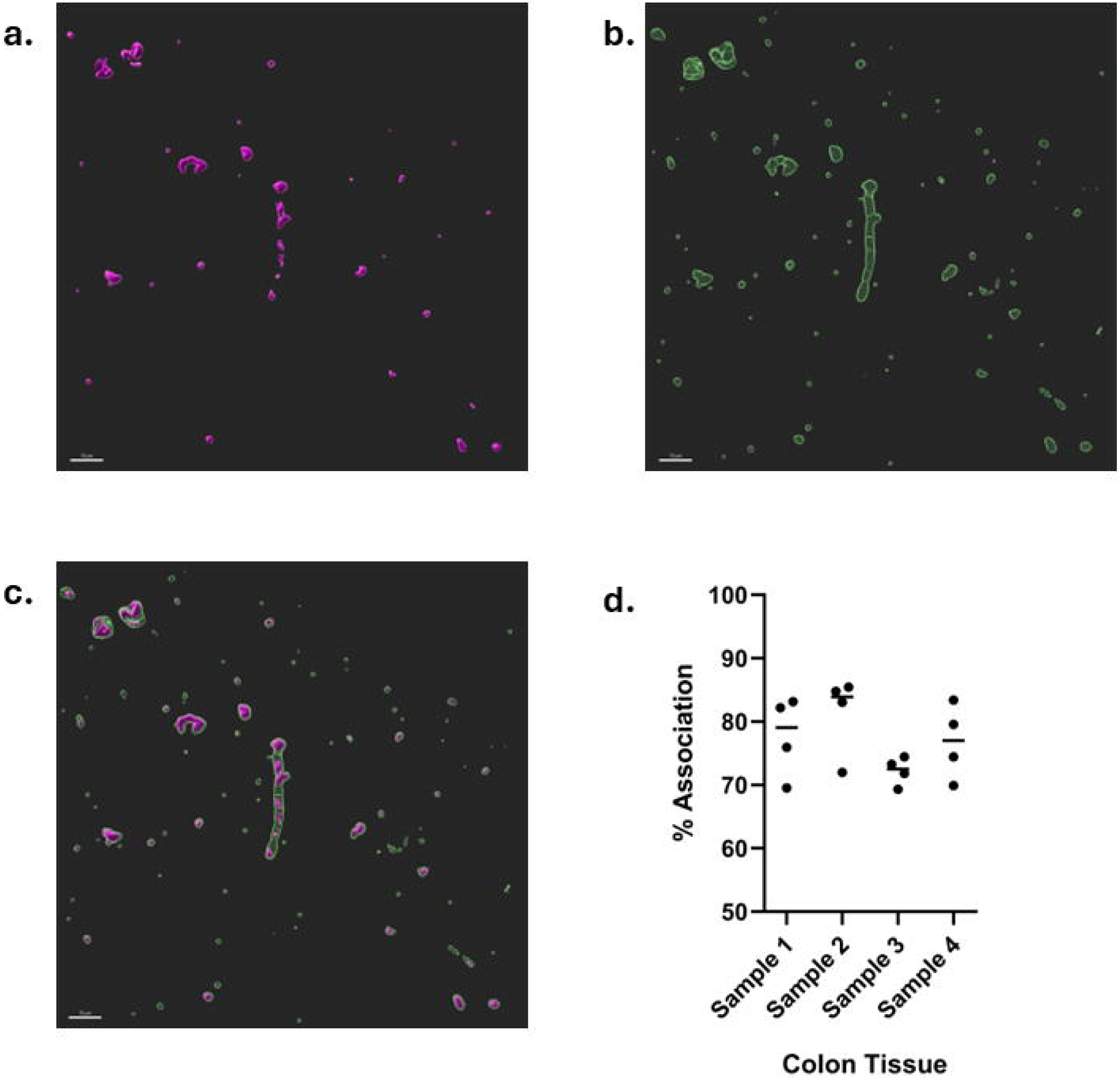
Laser confocal scanning microscopy evaluating GBS and *C. albicans* co-association in the colon of mouse pups. Representative rendered images of GBS and CA co-infected within the colon GBS only (**a**), CA only (**b**), and combined GBS and CA (**c**), and the relative percentage of GBS associated with CA, as opposed to GBS not spatially adjacent to CA was determined through quantitative analysis in Imaris (**d**). [green = CA, FITC; purple = GBS, anti-GBS IgG Alexa Fluor™ 568] Scale bar: 10 μm

### *C. albicans* induces changes in GBS gene expression

The expression of several GBS genes (*cylE, hvgA, srr2, fbsB,* and *spb1*) increased in response to the presence of *C. albicans* in co-culture (Fig. 5a). To determine whether these changes were dependent on the presence of live *C. albicans* or a secreted factor, we repeated these experiments using only *C. albicans* culture supernatant (Fig. 5b) and heat-killed *C. albicans* (Fig. 5c) incubated with GBS. In these conditions, *cylE*, *srr2*, and *spb1*, were similarly up-regulated. To determine whether *C. albicans* would exhibit an effect on GBS growth in the *in vitro* co-culture broth and pH values, a growth curve and pH assessment was conducted with no difference between monoculture and co-culture samples (Fig. S4).

**Figure 5.**
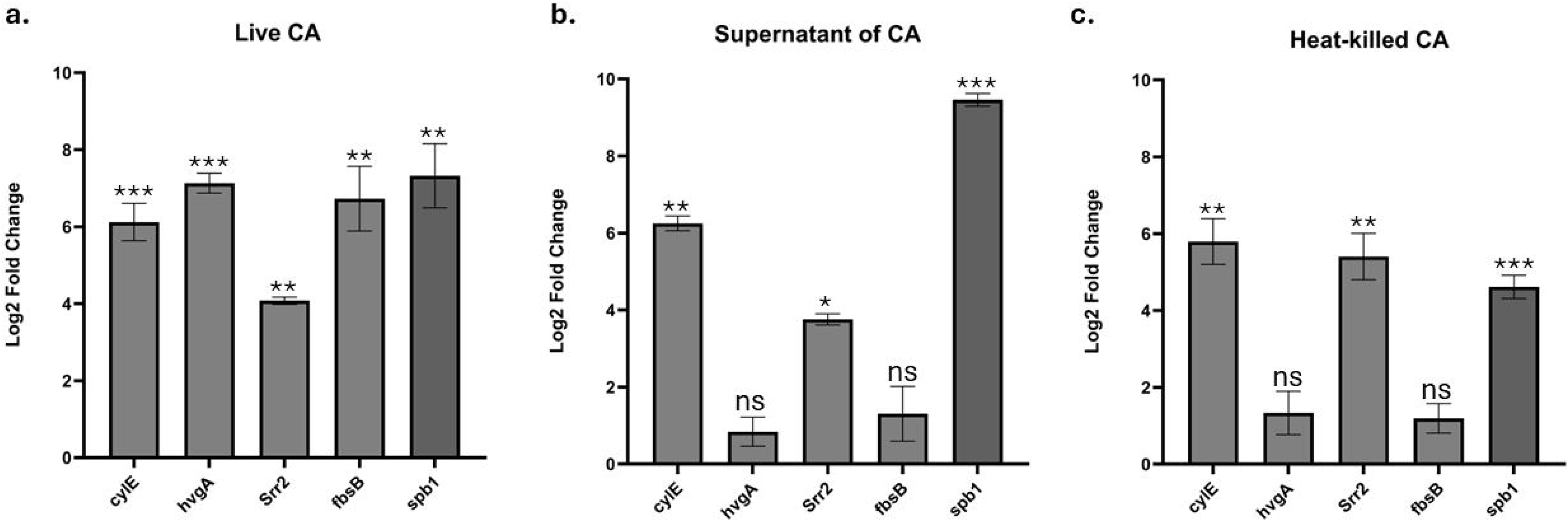
Candida albicans impact on GBS virulence. Relative gene expression of *cylE, hvgA, srr2, fbsB* and *spb1* normalized to *recA* in *in vitro* GBS co-cultured with either live CA (**a**), supernatant of CA (**b**), or heat-killed CA (**c**) for 4 hours compared to GBS monoculture baseline expression. [***p<0.0003, **p<0.008, *p<0.04, Welch’s t test]

### *C. albicans* presence increased GBS adherence to epithelial cells

Co-infection with *C. albicans* led to a marked increase in GBS adherence to T84 cells compared to those in GBS monoculture (Fig. 6a). Invasion of T84 cells remained unaffected by the presence of *C. albicans* (Fig. 6b). No significant differences in adhesion or invasion were detected in Caco-2 epithelial cells across conditions (data not shown). Cell viability was also evaluated to post infection incubation period (Fig. S5).

**Figure 6.**
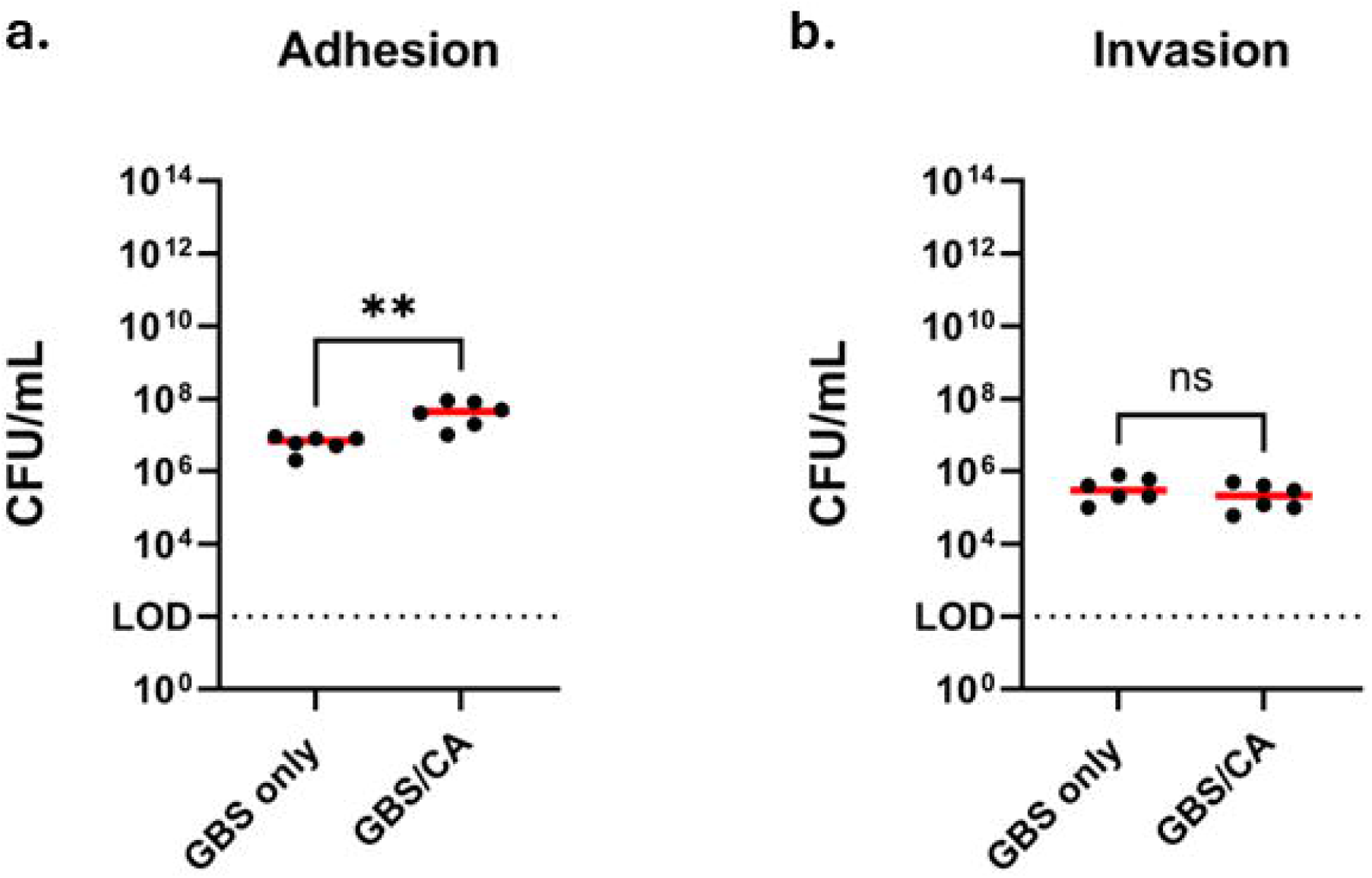
Adherence to and invasion of epithelial cells. Adhesion and invasion assays of T84 cells (**a-b**) at MOI 50 following 2-hour incubation with GBS monoinfection vs co-infection of GBS and CA. Limit of detection is 10^2^. [**P=0.002, ns=nonsignificant, Mann Whitney test]

## Discussion

In this study, we successfully established a novel reproducible neonatal model of gastrointestinal co-colonization with GBS and *C. albicans* and demonstrated that these fungal-bacterial interactions within the immature gut can significantly alter GBS pathogenic potential. Our findings show that the presence of *C. albicans* is associated with increased GBS intestinal burden and fluconazole treatment reduced GBS extraintestinal dissemination in co-colonized animals, suggesting that suppression of *C. albicans* may indirectly limit bacterial translocation. High-resolution *in vivo* imaging revealed close physical associations between GBS and *C. albicans*, similar to interactions previously described at other mucosal sites, including the bladder and vagina.^25,26^ In addition, we observed that *C. albicans* induced changes in the expression of GBS genes linked to pathogenicity, including hemolytic/cytolytic activity (*cylE*), adhesins (*hvgA* and *srr2*), and invasion of epithelial cells (*fbsB* and *spb1*).^30,31^ However, a limitation to our gene expression analyses performed *in vitro*, is that this does not fully capture the influence of the host microenvironment or the complexity of immune signaling that can significantly shape pathogen behavior. Additionally, our approach was limited to a few candidate genes. Future studies employing transcriptome-wide approaches or metatranscriptomics could provide a more comprehensive view, including insights into how *C. albicans* itself responds within this polymicrobial context. We also focused on a single strain of GBS. While this strain represents the sequence type of GBS most frequently responsible for LO disease in infants, extending these observations to additional clinically relevant isolates would strengthen the generalizability of our findings. Collectively these physical interactions and transcriptional adaptations may contribute to the increased persistence and dissemination of GBS in the neonatal gut.

These findings align with a growing body of literature demonstrating that cross-kingdom interactions between *Candida* species and bacteria may be critical factors that profoundly influence colonization, virulence, and host outcomes. Functional consequences of these interactions are evident in other coinfection models, where *C. albicans* and *S. aureus* co-inoculation in mice leads to rapid dissemination and increased mortality compared to monoinfections, suggesting synergistic virulence.^32–34^ Similarly, neonatal rat models demonstrate that co-colonization with *C. albicans* and *Staphylococcus epidermidis* results in impaired growth and increased morbidity, effects that can be mitigated with antifungal prophylaxis.^35^ At the structural level, dual-species biofilm studies reveal that *S. aureus* preferentially associates with *C. albicans* hyphae, forming dense polymicrobial communities that enhance persistence and resistance.^21,32,34^ Our imaging data suggest that similar physical associations may occur in the neonatal gut, providing a scaffold that facilitates GBS adherence, protection from host immunological defenses, or enhanced access to epithelial surfaces. Together, these findings underscore the bidirectional nature of fungal-bacterial interactions, where metabolic signals and physical associations shape pathogenic potential in the host.

The neonatal gut presents a uniquely permissive environment for these interactions. Compared to children and adults, infants (particularly preterm neonates) have an immature epithelial barrier, reduced mucus production, and limited tight junction integrity.^36,37^ These characteristics could enhance the probability of microbial translocation when pathogens are in high density. Additionally, the preterm infant immune system is vastly underdeveloped with fewer neutrophil populations, weaker Th1 and Th17 responses, and reduced production of antimicrobial peptides.^38,39^ In this context, the presence of *C. albicans* may further modulate immune signaling that could lead to skewing of the immature immune response in a way that could support GBS persistence within the gut. Additionally, the early-life (neonatal) microbiome exhibits low diversity and limited resistance to bacterial colonization, allowing GBS and *C. albicans* to grow and interact more easily and without the competitive constraints found in the mature gut environment.^40,41^

Several mechanistic pathways provide potential insight into how *C. albicans* alters GBS virulence and, potentially, dissemination. The hyphal structure could provide a physical relationship that encourages GBS adherence and possibly shields it from host immune clearance. While the relevance of this concept to GBS is not yet fully understood, evidence indicates a role of *C. albicans* hyphae in immune evasion. *C. albicans* hyphae accomplish immune evasion by escaping macrophages using the fungal toxin candidalysin to permeabilize the membrane of macrophages, thus facilitating *C. albicans* release while also activating Gasdermin D-mediated pyroptosis and ETosis host apoptosis pathways.^42^ This physical disruption of macrophages allows the fungal hyphae to evade phagocytic killing. GBS compromises the functionality of macrophages by regulating apoptosis and bactericidal activity, allowing it to persist intracellularly as an immune evasion tactic. GBS activates caspase-9 and caspase-3 and release of cytochrome c, altering the regulation of apoptosis pathways. GBS also uses the β-hemolysin cytotoxic *cylE* factor for bacterial release.^43,44^ GBS has also been noted to survive inside macrophages for extended periods of time and to evade immune detection by inhibiting reactive oxygen species and nitric oxide production.^45–47^ The interaction between *C. albicans* and GBS indicates pathogenic effects that can significantly compromise the immune defense of the host and contribute to systemic infection, particularly in susceptible individuals.

The relationship between the hyphal-associated adhesin Als3 from *C. albicans* and GBS adhesin factors have been previously described.^25,26^ Our *in vivo* imaging data revealing the physical interactions along with the increased gene expression related to the pathogenicity of GBS may provide mechanistic validation and suggest that GBS uses hyphae to evade host immune defense.

In this study, *C. albicans* was grown under continuous shaking overnight for murine experiments and, for gene expression experiments, it was grown stationary overnight. Previous studies have reported that stationary incubation allows for CO_2_ accumulation, and hypoxia subsequently promotes increased hyphal morphogenesis in *C. albicans*, whereas incubations with continuous shaking counteract these preferred hyphal conditions contributing to the maintenance of budding yeast morphology.^20,48^ This approach was selected to ensure morphological consistency between the *in vivo* and *in vitro* systems. In the LO disease neonatal mouse model, pups were colonized with *C. albicans* for 3 days prior to GBS challenge, a duration sufficient for the yeast to hyphal transition within the host environment. However, in the current study, the *in vitro* cultures were independently grown overnight followed by only a 4-hour co-incubation period, a timeframe insufficient for equivalent hyphal development. To approximate the morphogenetic state present during the *in vivo* colonization, *C. albicans* growth conditions were adjusted prior to *in vitro* co-incubation with GBS.

Other studies have shown how the structural features of *C. albicans* hyphae assist in reducing immune detection by inhibiting type 1 interferon (IFN-1) signaling, which is critical for immune defense against pathogens and important in early stages of innate immunity.^49^ GBS has also been reported to affect IFN-1 signaling in various ways^47,50^ and, in conjunction with the effects caused by *C. albicans*, neonatal immunity is further weakened and the downstream effects potentially amplified.

One important mechanism of cross-kingdom communication involves quorum sensing molecules (QSMs) such as farnesol, a fungal metabolite produced by *C. albicans* that regulates its yeast-to-hyphal transition. While farnesol limits fungal filamentation at high concentrations, bacteria have adapted to sense and respond to this signal. For instance, *Staphylococcus aureus* detects farnesol, which can disrupt bacterial membranes and reduce viability, whereas *Pseudomonas aeruginosa* downregulates its own QSM production upon sensing farnesol.^51–56^ Fluconazole exposure enhances *C. albicans* production of farnesol which represses the transition from yeast to hyphal growth, impairing the formation of mature biofilms and host tissue invasion.^57,58^ In the context of our fluconazole treated pups versus the *in vitro* heat-killed *C. albicans* and supernatant of *C. albicans* experiments, secreted metabolites such as farnesol and other volatile organic compounds could be released that can modulate bacterial motility and gene expression, highlighting the complexity of chemical signaling in mixed microbial communities.

Our observations of the effects of fluconazole treatment on co-colonized animals warrant careful consideration, especially since fluconazole is routinely used in neonatal intensive care units to lower the risk of invasive candidiasis in preterm infants. While fluconazole reduced GBS dissemination in our model, it is uncertain whether this effect would be consistent across different GBS strains or host species. The impact of the practice of fluconazole prophylaxis on the incidence of neonatal sepsis caused by enteric pathobionts such as GBS remains unclear and represents a critical area for future investigation.

Previous studies have revealed a critical need to investigate the interactions between *C. albicans* and GBS, particularly within vulnerable populations at the highest risk for invasive disease. Despite the clinical relevance of these organisms, there is a disparity in studies focusing on these specific interactions within the preterm infant gut whose immature intestinal and immune barriers predispose them to polymicrobial pathogenesis. To address this gap, we successfully created a reproducible mouse model of GBS and *C. albicans* intestinal co-colonization revealing increased GBS in the murine neonatal gut when *C. albicans* was present. Mechanistic analysis of this relationship revealed that *C. albicans* impacts gene expression of GBS associated with virulence and host invasion. Defining the specific interactions of polymicrobial dynamics within the human infant gut such as the synergistic behaviors of GBS and *C. albicans* that can impact and lead to neonatal sepsis is critical for advancing early diagnostic approaches as well as developing targeted interventions to prevent profound neurological impairment and mortality.

## Data availability

The data generated and analyzed during this study are available from the corresponding author on reasonable request.

## Acknowledgements

We would like to acknowledge Jane Carver, PhD, MS, MPH for her input in reviewing this manuscript.

## Author Contributions

TMR conceived the project. AKL, ANP, KD, NJE, RH, CZA, SED, and TMR performed experiments and analyzed data. AKL, ANP, KD, OMW, ON, LJD, SED, and TMR contributed to experimental design, interpreting data, and drafting the manuscript. All authors contributed to the article and approved the submitted version.

## Ethics Approval

All experiments were approved by and performed in accordance with the University of South Florida Institutional Animal Care and Use Committee.

