## Supplemental Figures and Figure Legends for "Microbial Crosstalk in the Neonatal Gut: *Candida albicans* Impacts Group B *Streptococcus* Pathogenicity"

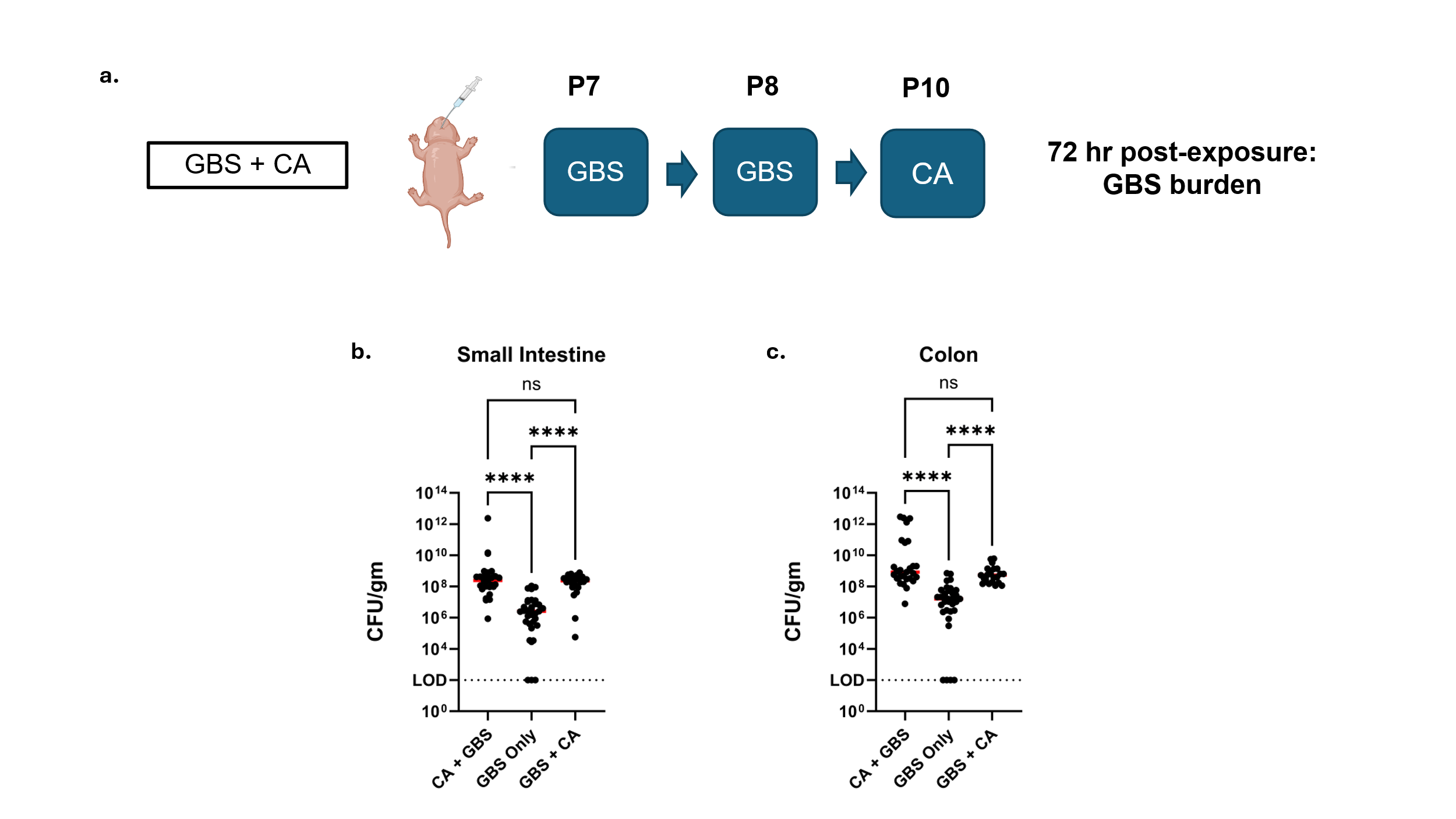

Figure S1. **GBS burden from reverse infection model**. GBS and CA reverse co-infection model (**a**) by infecting with 2 doses of GBS on day of life 7 and 8 final dose of 10^7^ CFU daily followed by infection with CA with a dose of 5 X 10^6^ CFU on day of life 10. GBS burden in the small intestine (GBS + CA median CFU = 1.8 X 10^8^ CFU/gm tissue versus GBS monoinfection median CFU = 4 X 10^6^ CFU/gm tissue) (**b**) and colon (GBS + CA median CFU = 6.3 X 10^8^ CFU/gm tissue versus GBS monoinfection median CFU = 1.6 X 10^7^ CFU/gm tissue) (**c**) was evaluated using serial dilutions. Statistical analyses were conducted using Graphpad Prism v11.0.2 [****P<0.0001, ns=nonsignificant, Kruskal-Wallis with Dunn’s Correction, LOD=limit of detection=10^2^].

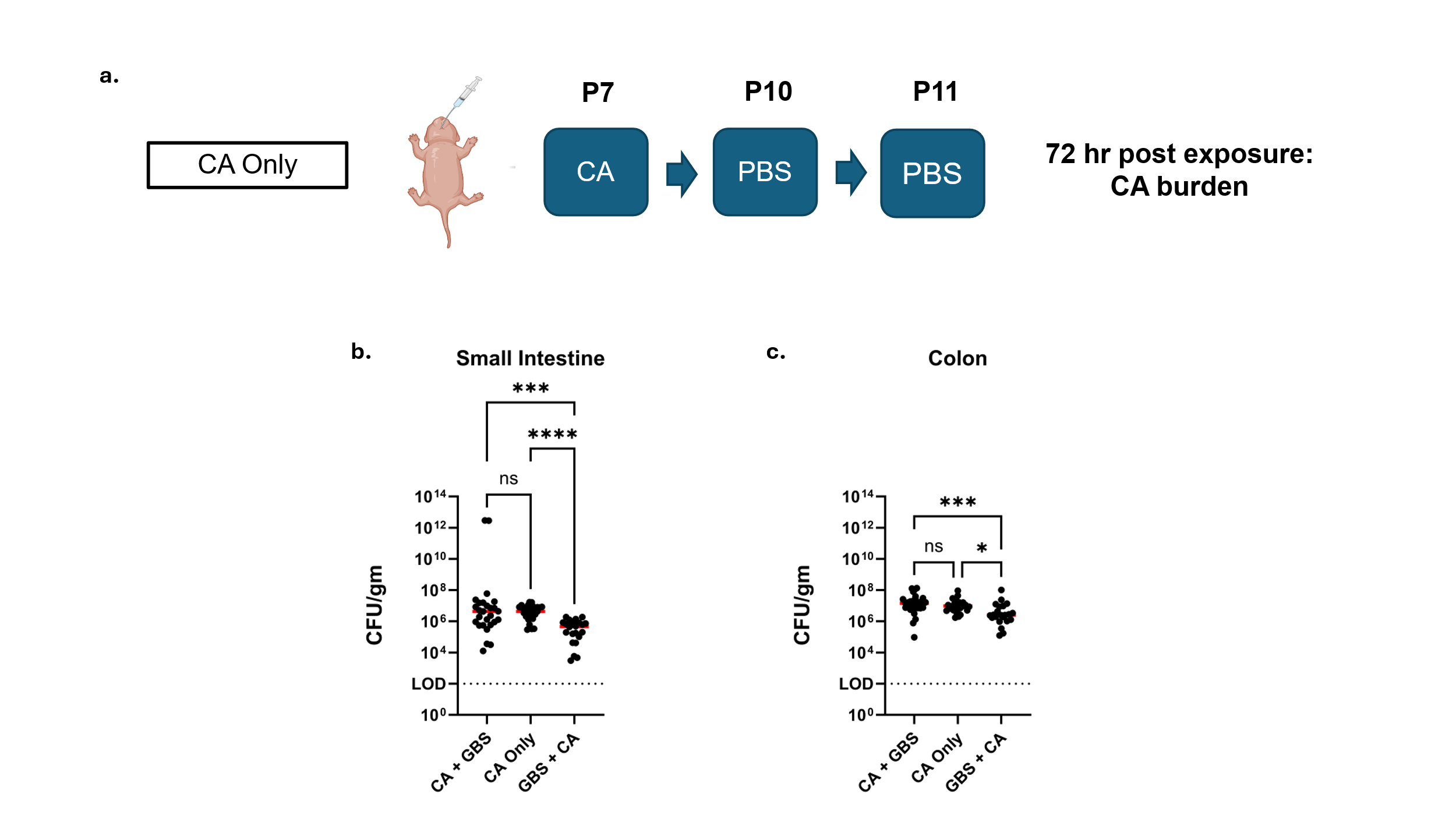

Figure S2. ***C. albicans* monoinfection model and burden.** CA monoinfection model (**a**) CA was administered with a dose of 5 X 10^6^ CFU on day of life 7 and PBS on day of life 10 and 11 instead of GBS. CA burden was evaluated by completing serial dilutions 72 hours post exposure of the small intestine (**b**) and colon (**c**). Statistical analyses were conducted using Graphpad Prism v11.0.2 [****P<0.0001, ***P<0.0005, *P<0.02, ns=nonsignificant, Kruskal-Wallis with Dunn’s Correction, LOD=limit of detection=10^2^].

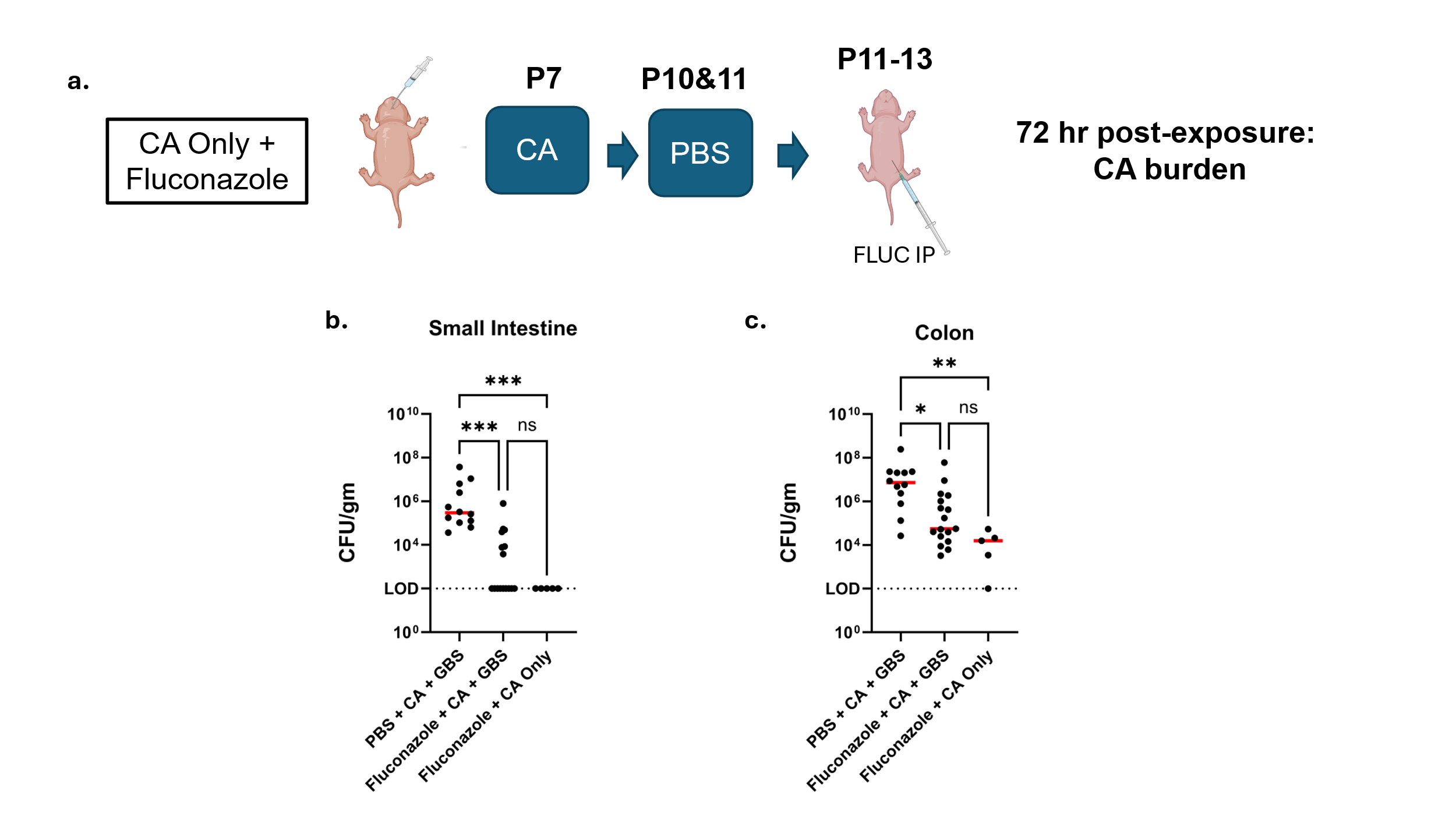

Figure S3. ***C. albicans* monoinfection and Fluconazole treatment model**. Monoinfection model of *Candida albicans* with a dose of 5 X 10^6^ CFU on day of life 7 and PBS on day of life 10 and 11 with Fluconazole (Sigma F8929, 5mg/kg daily, intraperitoneal) or PBS (**a**). CA burden in the small intestine (**b**) and colon (**c**) was evaluated using serial dilutions. Statistical analyses were conducted using Graphpad Prism v11.0.2 [***P<0.0009, **P<0.002, *P<0.03, ns=nonsignificant, Kruskal-Wallis with Dunn’s Correction, LOD=limit of detection=10^2^].

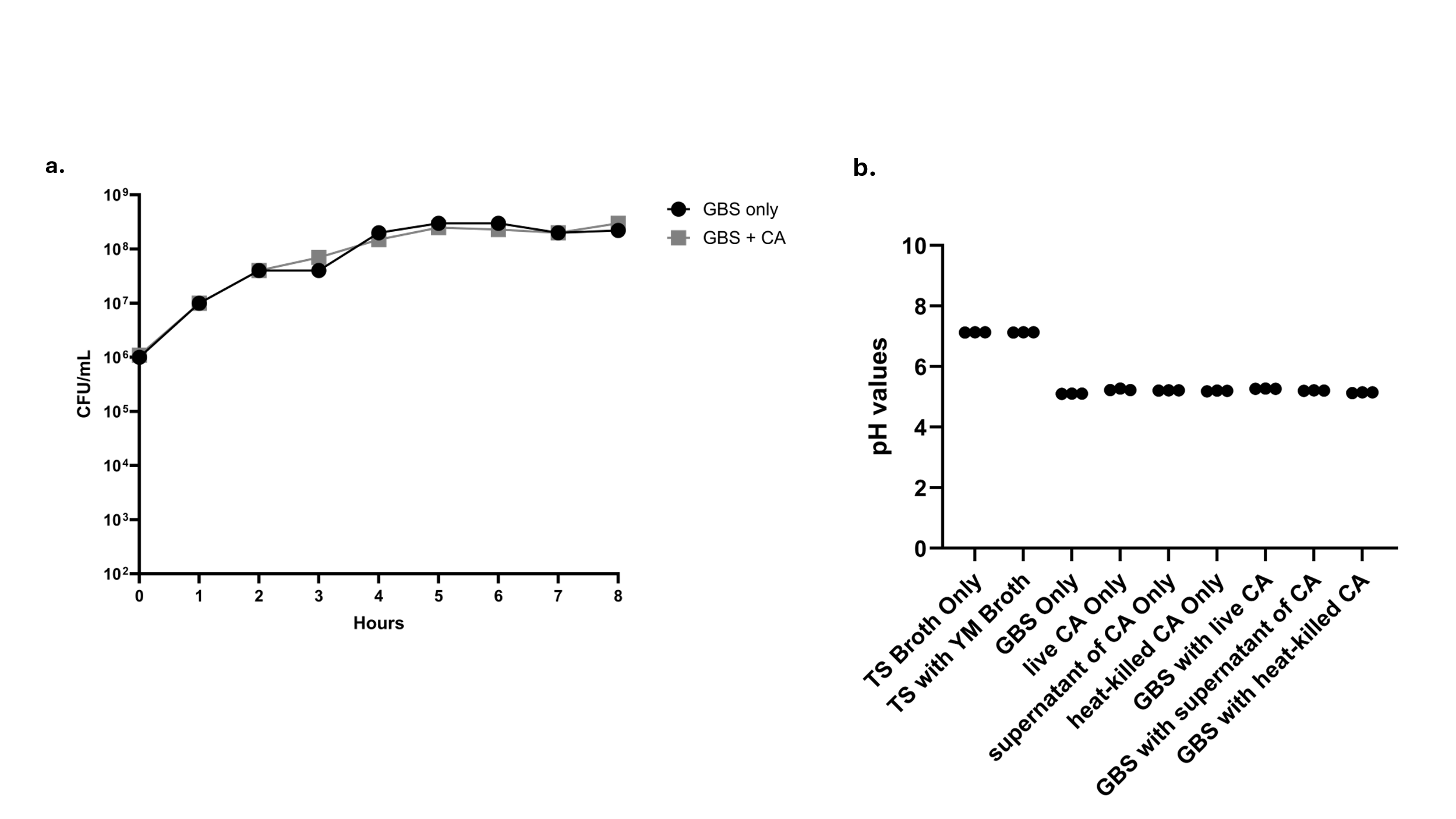

Figure S4. **Growth curve and pH values of GBS and *C. albicans* conditions**. GBS and *C. albicans* were cultivated in liquid broth overnight as described above, centrifuged and resuspended in 5 mL TS broth for GBS and 1 mL resuspension for *C. albicans*. Subcultures containing a 1:100 dilution of GBS monoculture and 1:100 dilution of GBS-*C. albicans* co-culture in TS broth were incubated in a 37°C water bath for 8 hours. Serial dilutions were plated every hour on CHROMagar^TM^ Strep B plates and CHROMagar^TM^ Candida plates to the 9^th^. CFU was evaluated for GBS the next day and CFU for *C. albicans* CFU was completed 2 days after plating (**a**). For pH assessment, GBS and *C. albicans* were cultivated in liquid broth overnight, centrifuged and resuspended as described previously. For co-culture of GBS with supernatant of *C. albicans*, following centrifugation, 4 mLs of supernatant were collected for coincubation. To heat-kill *C. albicans*, culture was pelleted, resuspended in YM broth, placed in the oven at 90°C for 1 hour and allowed to cool before co-incubating with GBS. All *in vitro* samples were incubated for 4 hours in a 37°C water bath and pH recorded with the accumet AB315 pH meter (**b**).

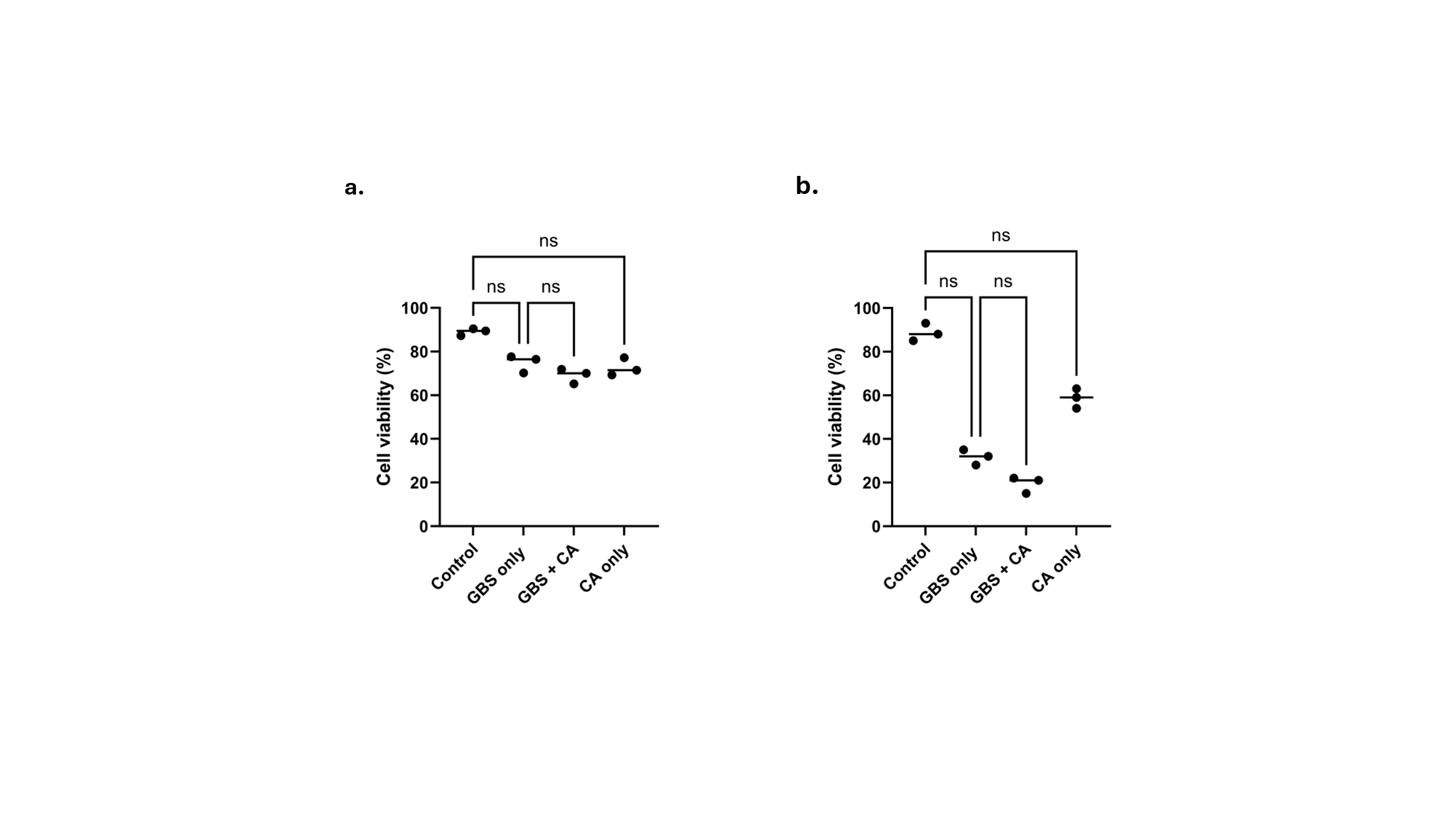

Figure S5. **Cytotoxicity assay of intestinal epithelial cells**. The trypan blue exclusion test to determine the viability of Caco-2 and T-84 cells as previously described.^1^ Briefly, the cell suspension was mixed with trypan blue dye, and the percentage of viable cells was determined by dividing the total number of viable cells per milliliter (unstained) by the total number of cells per milliliter (unstained + stained). Cell viability was evaluated of Caco-2 (**a**) and T84 cells (**b**) exposed to GBS and CA at MOI 50. Experiments were conducted in triplicate, at least, per group. Statistical analyses were conducted using Graphpad Prism v11.0.2 [ns=nonsignificant, Kruskal-Wallis test with Dunn’s Correction].

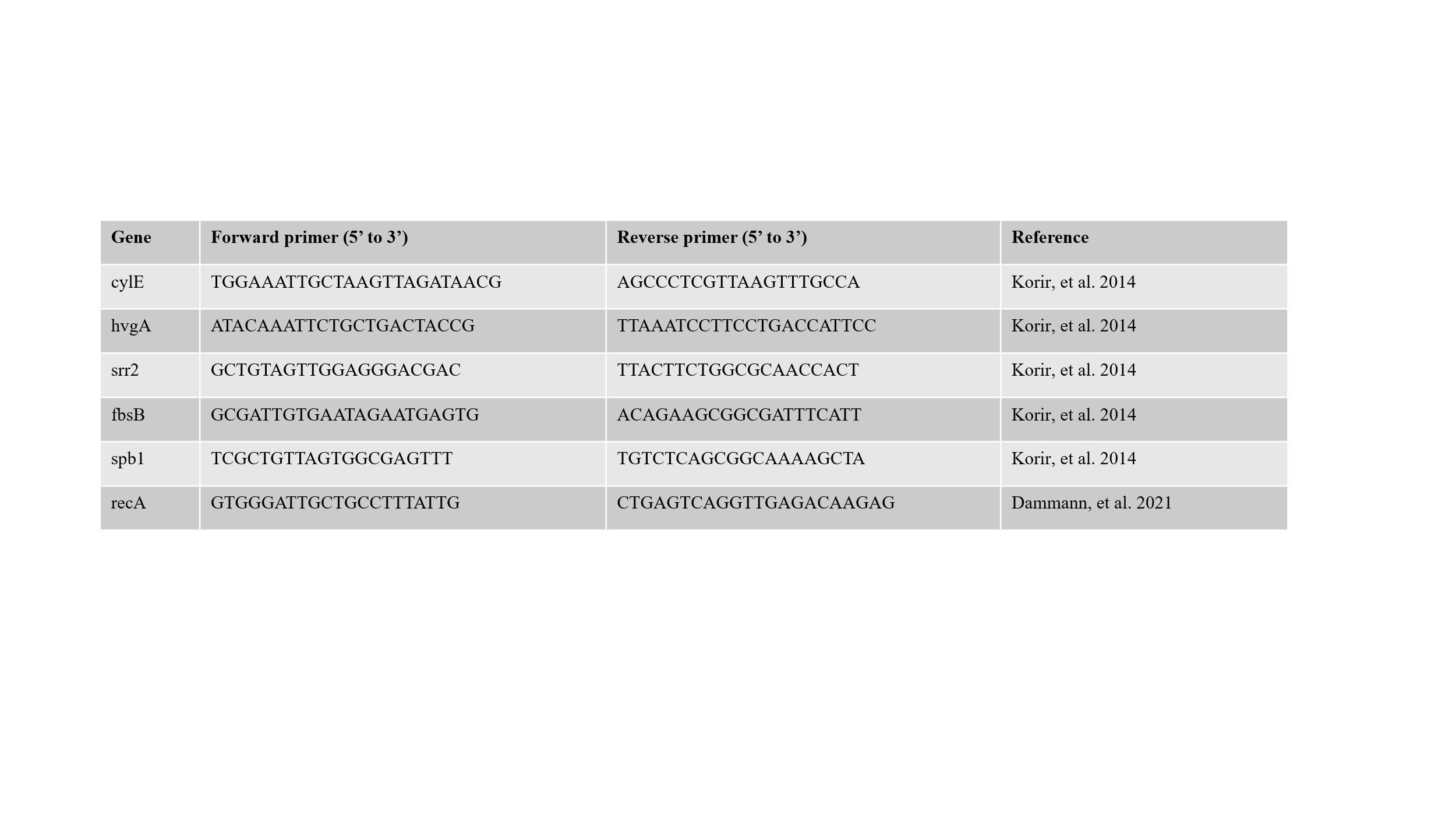

Table S1. **Primers for gene expression.** Previously published primers used in this study for gene expression of GBS.
